# Dithionite quenching of NBD-labeled lipids reveals artificial lipid droplet purity and neutral lipid surface accessibility

**DOI:** 10.64898/2026.07.07.737042

**Authors:** Jiyao Chai, Lingshuang Wu, Yong Mi Choi, Shujuan Gao, Daniel Canals, Abdou Rachid Thiam, Erwin London, Michael V. Airola

## Abstract

Artificial lipid droplets (aLDs) provide a controllable platform for studying lipid biochemistry, but their use is limited by contamination with other membrane structures and the lack of quantitative methods to assess sample purity. Here, we establish dithionite quenching of NBD-labeled lipids as a simple approach to evaluate aLD purity. The approach relies on dithionite’s ability to selectively quench NBD fluorophores exposed in the phospholipid monolayer of aLDs and in the outer leaflet of liposome bilayers, but not those protected within the inner leaflet of liposome bilayers. Consistent with liposome contamination, bulk aLD preparations exhibit incomplete quenching, which can be separated by sucrose gradient centrifugation into liposome-like and droplet-enriched populations based on quenching behavior. Guided by this assay, sonication conditions were optimized to increase aLD purity and reduce liposome contamination. A biotin–streptavidin immobilization strategy further enabled stable imaging of individual aLDs. Finally, we applied dithionite quenching to probe the accessibility of neutral lipids within aLDs. This revealed hydrophobicity-dependent quenching kinetics of neutral lipids, with less hydrophobic diacylglycerols showing greater surface exposure within aLDs than more hydrophobic triacylglycerols and cholesterol esters. Taken together, these establish dithionite quenching of NBD-labeled lipids as a simple quantitative method for assessing aLD purity and demonstrate its utility for studying lipid accessibility.

## INTRODUCTION

Lipid droplets (LDs) are ubiquitous organelles that store neutral lipids, primarily triacylglycerol (TAG) and cholesteryl esters, in a hydrophobic core surrounded by a phospholipid monolayer and associated proteins (1–5). LDs are now recognized as dynamic hubs for lipid metabolism, stress responses, and organelle communication, rather than inert storage sites (6–12). Central to these dynamic functions are lipid-metabolizing enzymes at the LD surface, including lipases that hydrolyze stored neutral lipids and acyltransferases that remodel the LD lipid composition (13–19). Because many LD-associated enzymes act at the monolayer interface while accessing substrates in the neutral lipid core, mechanistic studies of these enzymes require model systems in which lipid composition, surface properties, and substrate accessibility can be precisely controlled (20–26).

In vitro membrane models such as liposomes, which are phospholipid bilayer vesicles that can be unilamellar or multilamellar, are widely used for lipid enzymology; however, the bilayer architecture of liposomes fundamentally differs from the LD phospholipid monolayer that surrounds a neutral lipid oil core (1, 4, 5, 9). These architectural differences are critical because lipid packing, curvature, and leaflet topology can alter enzyme recruitment and substrate access (10–12, 15, 27). Thus, artificial lipid droplets (aLDs), which are oil-in-water emulsion droplets consisting of a neutral-lipid oil core enclosed by a phospholipid monolayer, have emerged as useful in vitro platforms for recapitulating LD architecture and supporting quantitative biochemical and biophysical assays under defined conditions (17, 18, 23, 24, 28–30).

Recent work has used aLD systems to study protein recruitment to LDs, droplet phase behavior, and enzyme-specific reactivity (23, 25, 29–32). In these approaches, fluorescence microscopy and electron microscopy are commonly used to confirm droplet morphology and provide evidence for monolayer-coated oil droplets (33, 34). However, morphology-based characterization does not quantify contaminating bilayer structures such as liposomes that can form during emulsification and sonication and may influence bulk biochemical readouts (24, 35). Reducing the production of non-lipid droplet vesicles is especially critical for enzymatic (e.g. lipid droplet lipase assays) (36) or lipid transfer assays (37) whose interpretation is complicated by contamination with non-lipid droplet vesicles. A rapid, quantitative method to assess aLD purity is therefore critically needed to optimize aLD preparations and ensure interpretability in biochemical assays.

To address this need, sodium dithionite quenching of NBD-labeled lipids provides a well-established chemical strategy to report fluorophore accessibility and membrane leaflet topology (26, 33, 34, 38, 39). Because dithionite selectively reduces solvent-exposed NBD groups, thereby quenching their fluorescence, differences in quenching efficiency can distinguish monolayer-coated droplets from bilayer structures and provide kinetic information about probe accessibility (34, 40). Here, we combine NBD-labeled lipids with dithionite quenching to detect and quantify liposome contamination in bulk aLD preparations. We subsequently apply this method to guide optimization of aLD preparations to reduce contaminating membrane structures and improve lipid partitioning between droplet and liposome fractions. Using this optimized system, we demonstrate that bulk aLD preparations support functional biochemical applications, including enzyme activity assays and assessment of neutral lipid accessibility at the LD surface.

## RESULTS

We sought to establish a method to assess the purity of artificial lipid droplets (aLDs) that did not involve microscopy, suitable for aLD characterization (17, 18, 24, 30), but that may fail to identify alternative membranous structures not visible by image analysis. aLDs were initially produced using a series of steps **(Supplemental Fig. 1A)** that were adapted from prior methods (24, 30) for in vitro enzyme assays (36). Fluorescence microscopy verified the presence of aLDs with a canonical architecture of a phospholipid (PL) monolayer surrounding an oil core of triglycerides **(Fig. 1A)**.

**Figure 1.**
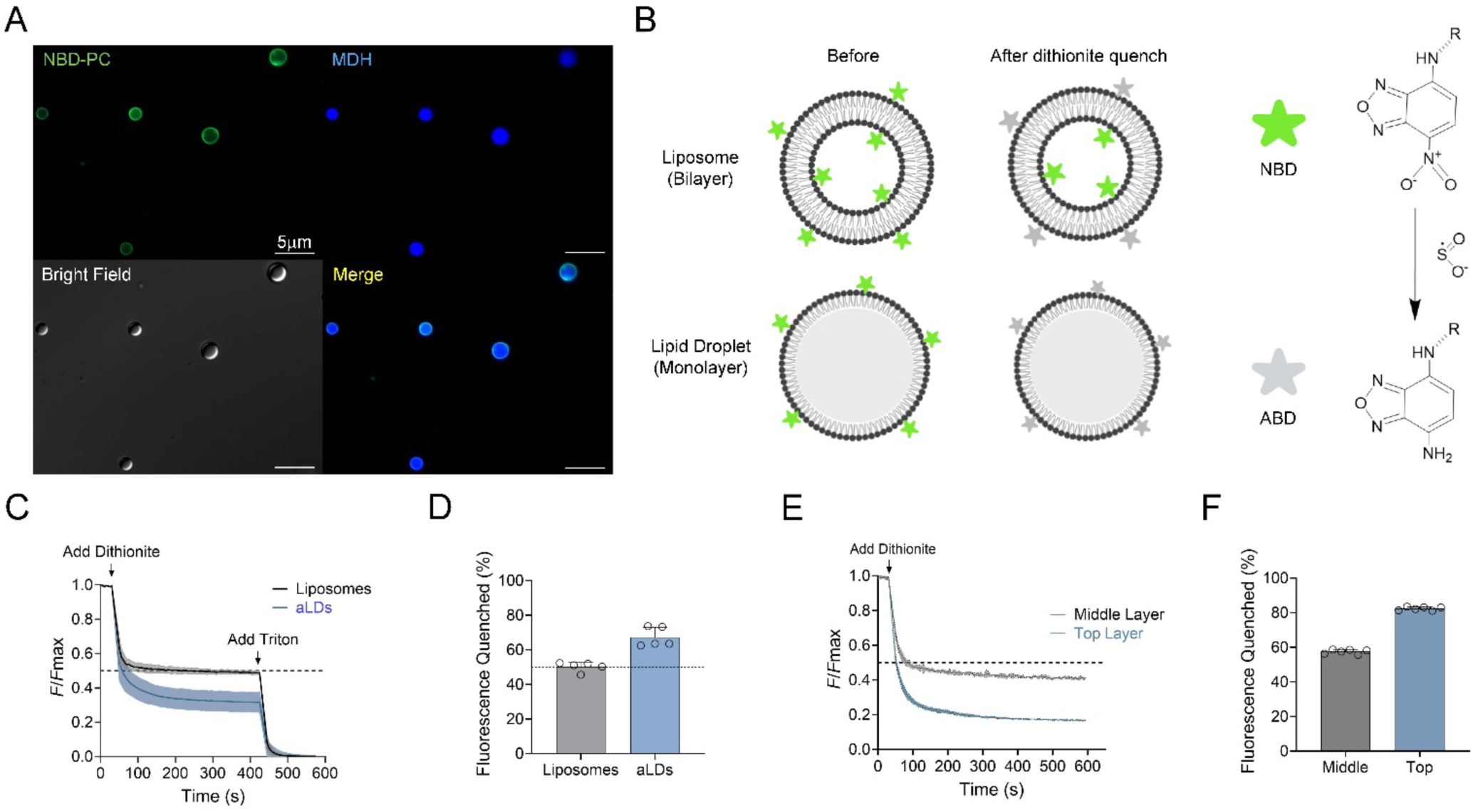
Dithionite quenching reveals liposome contamination in aLDs preparations. **(A)** Fluorescent microscopy images of artificial lipid droplets (aLDs) prepared with NBD-phosphatidylcholine (NBD-PC) to mark the phospholipid monolayer (green) and stained with Monodansylpentane (MDH) to mark the oil core (blue). **(B)** A diagram of NBD-phosphatidylethanolamine (NBD-PE) incorporated into either liposomes or aLDs before and after quenching with dithionite. **(C)** Normalized fluorescence intensity of liposomes or aLDs containing 1 mol% NBD-PE over time following the addition of sodium dithionite. Subsequent addition of Triton X-100 disrupts all membrane structures, which confirms the remaining fluorescence originates from protected NBD molecules. **(D)** Quantification of the quenching efficiency of liposomes and aLDs in **C** after 350 s. Mean ± SD is shown for each bar graph (n = 5). **(E)** Normalized fluorescence intensity after addition of dithionite of the top and middle (0–30% sucrose) layers collected after centrifugation of aLD preparations containing 1 mol% NBD-PE. **(F)** Quantification of the quenching efficiency of the top and middle layers in **E** after 350 s. Mean ± SD is shown for each bar graph (n = 5).

### Dithionite quenching reveals liposome contamination in aLD preparations

To assess aLD purity, we developed an approach using dithionite quenching of NBD-labeled phosphatidylethanolamine (PE) where NBD is on the headgroup **(Fig. 1B)**. This approach aimed to take advantage of the different quenching efficiencies of NBD-labeled PLs when incorporated into aLDs or liposomes **(Fig. 1B)**. As previously characterized (39), large unilamellar liposomes with NBD-PLs give rise to 50% quenching due to the inaccessibility of the NBD-groups to dithionite on the inner leaflet of the membrane bilayer. We reasoned that pure aLDs would alternatively give rise to 100% quenching, as previously observed by microscopy for single aLDs (24), due to the complete solvent exposure of the NBD-PLs in the aLD monolayer. A readout of contamination in aLD preparations would thus be interpreted as an intermediate degree of quenching between 50-100%, with aLD purity proportional to the degree of NBD-dithionite quenching.

As a control, unilamellar liposomes containing 1 mol% NBD-PE were prepared and showed ∼50% fluorescence quenching upon addition of dithionite **(Fig. 1C, D)**. This could be converted to 100% quenching after the addition of Triton X-100 detergent, which disrupts all membranous structures **(Fig. 1C)**. In comparison, aLDs prepared using our initial protocol displayed ∼65% quenching **(Fig. 1C, D)**. This suggested that our aLD preparations also contained a significant fraction of liposomes.

To confirm the presence of liposomes in aLD preparations, we used sucrose gradient centrifugation to separate aLDs from liposomes, followed by the separate collection of each layer and subsequent dithionite quenching. aLDs were generated in the presence of 15% sucrose, which would be encapsulated in liposomes but not aLDs. This would allow separation of liposomes from aLDs by sucrose gradient centrifugation **(Supplemental Figure 1B)**.

After centrifugation, there were clear layers of lipid in the middle layer, where sucrose-laden liposomes should reside, and in the top layer, where less dense aLDs should float. The middle layer displayed a ∼50% quenching extent, consistent with that of unilamellar liposomes **(Fig. 1E, F)**. In comparison, the top layer, which contained the semi-purified aLDs, had an improved quenching extent of ∼80% **(Fig. 1E, F)**. Taken together, we concluded that our protocol for generating aLDs also produced contaminating liposomes, and that dithionite NBD-PL quenching provides a rapid method to evaluate the purity of aLD preparations.

### Protocol optimization to increase aLD purity

We next sought to optimize our protocol to reduce liposome contamination, improve aLD purity, and eliminate the need for centrifugation. Given that our initial protocol involved several steps and variables **(Supplemental Fig. 1A)**, we independently changed each variable and assessed aLD purity using dithionite quenching.

We found two major variables that improved aLD purity. Both variables involved the tip sonication step. The first variable was the amplitude of tip sonication. Increasing the sonication amplitude from 25% to 40% improved quenching from 50 to 60% when tip sonication was carried out at 25°C **(Fig. 2A)**. A second important factor was the temperature of the water bath during tip sonication. By raising the temperature from 25°C to 50°C during tip sonication, the dithionite quenching increased from 60% to ∼75% **(Fig. 2B)**. All other variables tested, including variations in PL acyl-chains, triolein concentration, and PL concentration, did not result in any statistically significant improvement of aLD purity as assessed by dithionite quenching **(Fig. 2C-F)**. Although ultrasonic power (amplitude) is the primary factor driving emulsification efficiency and droplet size reduction, variations in sonicator probe tip diameter can influence the energy provided to the mixture (41). Therefore, our optimized parameters may not directly translate to other models but can be easily optimized for each system for the NBD-dithionite quenching assay.

**Figure 2.**
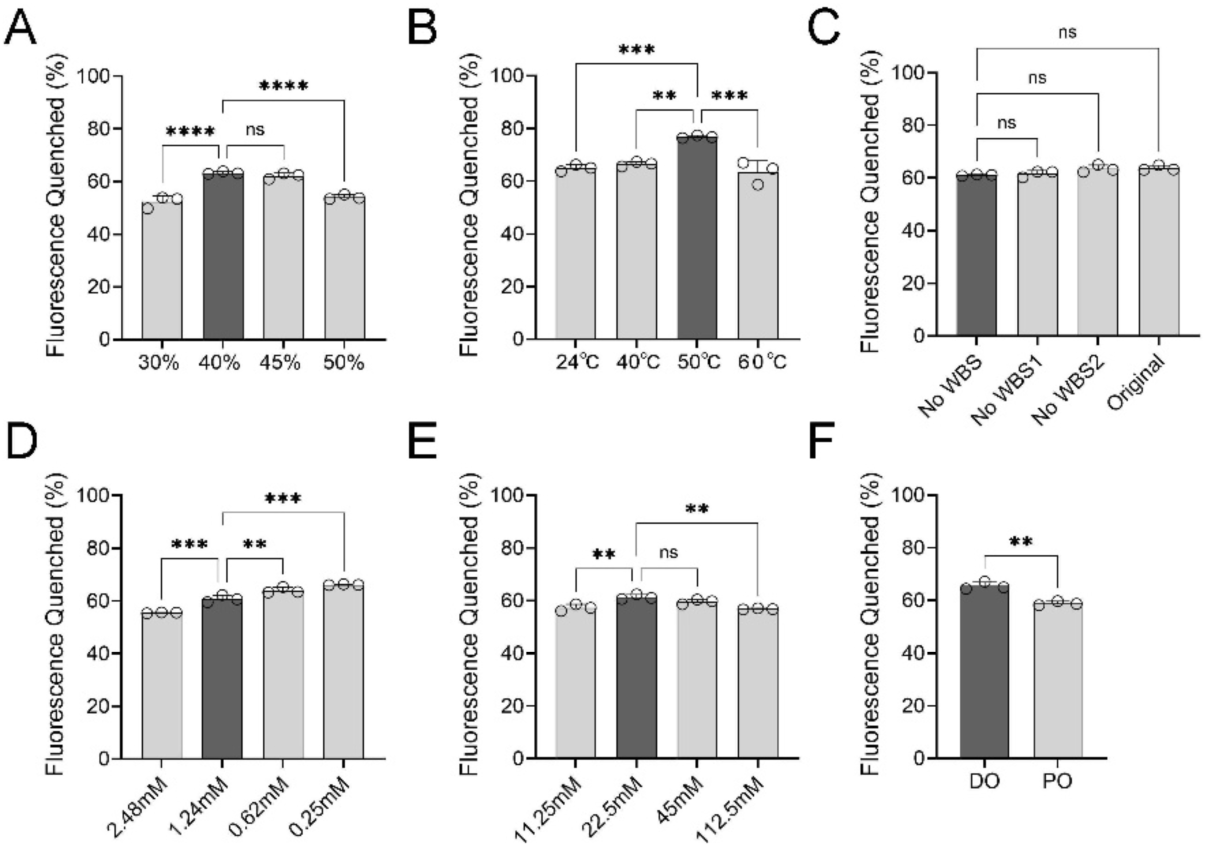
Dependence of aLD purity on the tip sonication parameters. (A-F) Quantification of the quenching efficiency of aLDs after 350 s as a function of **(A)** tip sonication amplitude, **(B)** tip sonication temperature, **(C)** protocol water bath sonication steps (WBS: water bath sonication), **(D)** phospholipid concentration (keep TAG concentration 22.5mM), **(E)** triolein concentration (keep PLs concentration 1.24mM), and **(F)** phospholipids acyl-chain saturation. Mean ± SD is shown for each bar graph (n = 3), significance is calculated by One-Way ANOVA. The dark grey bar in each panel represents the condition selected for each variable in the optimized aLD protocol.

We also assessed which steps were critical to generate aLDs in our protocol. We found that the tip sonication step was the primary determinant of aLD purity, and that leaving out either or both of our initial water bath sonication steps **(Supplementary Fig. 1A)** did not affect the overall purity **(Fig. 2C)**. After optimization, our protocol could generate aLDs with 75% dithionite quenching, which was comparable to purification by centrifugation. We concluded that aLDs of similar purity could be prepared without having to undergo any centrifugation steps.

### Biotin–streptavidin immobilization enables stable visualization of artificial lipid droplets

During fluorescence microscopy of artificial lipid droplets (aLDs), stable imaging requires minimizing droplet deformation and movement. It is critical to pretreat the coverslip and slide with 10% (w/v) BSA to prevent droplet adhesion and allow aLDs to maintain their spherical structure (18). Despite this treatment, aLDs were not stably anchored to the imaging surface, which occasionally caused positional shifts during z-stack acquisition. To address this issue, we tested a biotin–streptavidin coupling strategy to immobilize aLDs on the coverslip for microscopy.

Biotin–streptavidin interactions have been widely used to tether or functionalize lipid assemblies with high affinity and minimal disruption to membrane structure (42, 43). Biotinylated phospholipids, such as biotin-PE, enable streptavidin-mediated attachment of lipid droplets to solid surfaces or other biomolecules (44, 45). We adapted this approach by replacing the standard BSA coating on the coverslip with biotin-labeled BSA. After washing and incubation with streptavidin, a binding layer was formed that could capture biotin-PE–containing aLDs through biotin–streptavidin linkage **(Fig. 3A)**.

**Figure 3.**
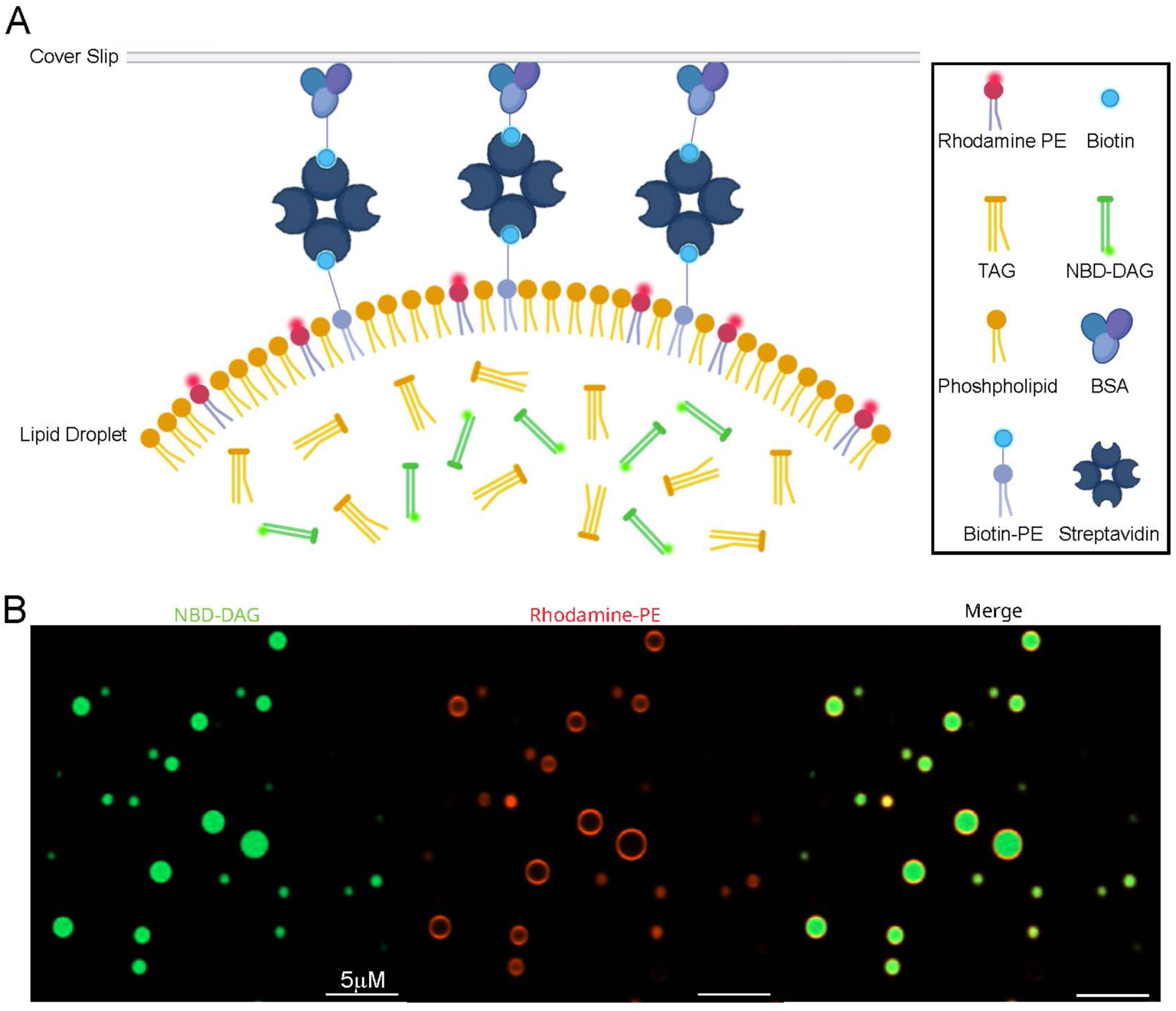
Biotin–streptavidin immobilization enables stable visualization of artificial lipid droplets. **(A)** Diagram of the biotin-streptavidin immobilization system to immobilize aLDs for stable microscopy visualization. Streptavidin forms a bridge between the BSA-biotin coating on the glass coverslips and biotin-PE that is incorporated into the PL monolayer of aLDs. **(B)** Confocal microscopy images of biotin-streptavidin immobilized aLDs prepared with NBD-DAG (green) and rhodamine-PE (red). Scale bar = 5 μm

Airyscan confocal imaging of immobilized aLDs revealed a clear ring-shaped Rhodamine-PE signal outlining the droplet membrane and an NBD-DAG fluorescence within the hydrophobic core. Importantly, the droplet positions remained stable throughout imaging **(Fig. 3B)**. The immobilized preparations could be sealed and stored for up to a week, thus providing a reproducible and convenient platform for fluorescence-based analysis of aLDs.

### NBD-PC provides superior dithionite-quenching sensitivity for assessing aLD purity

Fluorescence microscopy observation of NBD-PE **(Fig. 4A)** labeled aLDs before and after dithionite addition showed that the surface fluorescence was not completely quenched **(Fig. 4B)**. This suggested that NBD-PE may not fully reflect the accessibility of phospholipids on the aLD surface or the actual purity of aLD preparations. To test whether another fluorescent lipid could improve detection sensitivity, NBD-PC **(Fig. 4A)** was examined. The NBD group in NBD-PC is attached to the acyl chain rather than the headgroup. We reasoned that maintaining the native headgroup of PC and attaching the NBD group to the acyl chain may lead to a more exposed fluorophore, given that NBD groups in the acyl chains have previously been shown to be solvent exposed in liposomes (26, 33).

**Figure 4.**
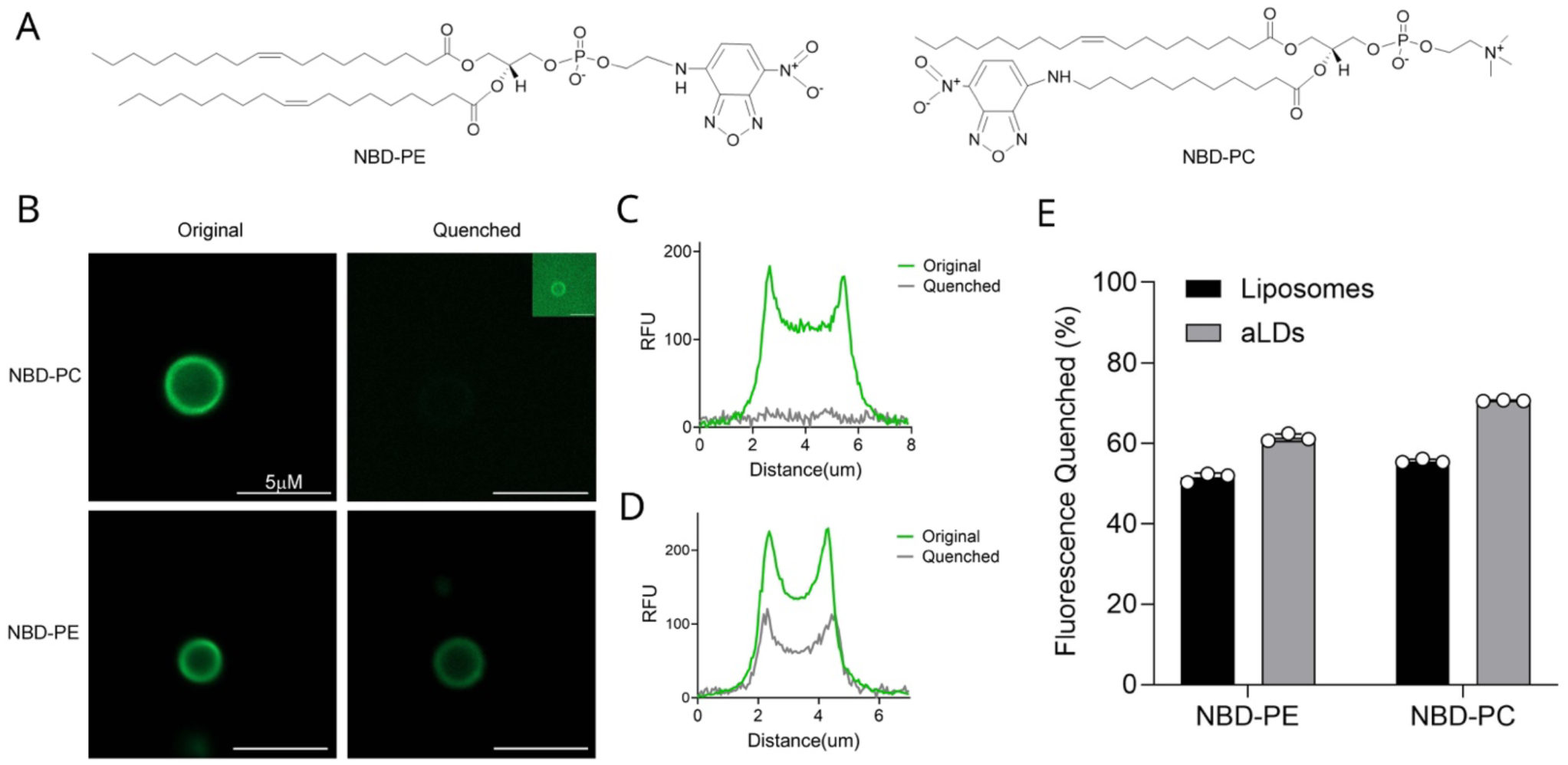
NBD-PC provides superior dithionite-quenching efficiency to assess aLD purity. **(A)** Chemical structures of NBD-PE and NBD-PC showing the different position of NBD on the headgroup of NBD-PE versus the acyl-chain of NBD-PC. **(B)** Fluorescence microscopy images of NBD-PC or NBD-PE labeled aLDs before and after dithionite quenching. Scale bar=5 um. The fluorescence intensity was adjusted in the inset (top right) to show the very low residual fluorescence of NBD-PC after dithionite quenching. **(C-D)** Fluorescence intensity line function comparison before and after dithionite quenching of (C) NBD-PC and (D) NBD-PE labeled aLDs. **(E)** Comparison of the fluorescence dithionite quenching of liposomes and aLDs prepared with either NBD-PE or NBD-PC after 350 s. Mean ± SD is shown for each bar graph (n = 3).

As an initial experiment, aLDs containing either NBD-PE or NBD-PC were prepared using identical protocols and imaged before and after dithionite treatment **(Fig. 4B)**. Both NBD-labeled phospholipids located predominantly at the surface of the LD in a fashion similar to Rhodamine-PE (**Fig. 4B**). Liposome samples labeled with the same probes were included as controls. aLDs labeled with NBD-PC showed a near complete loss of fluorescence after dithionite quenching **(Fig. 4B, C)**, while NBD-PE–labeled aLDs retained a weak signal **(Fig. 4B, D)**. Liposomes containing NBD-PC displayed ∼55% quenching **(Fig. 4E)**, consistent with their bilayer structure, where half of the probes face the inner leaflet.

When the quenching levels in aLDs were compared, NBD-PC labeling resulted in 72% fluorescence loss, whereas NBD-PE reached only 64% under identical conditions **(Fig. 4E)**. These results indicate that NBD-PC is more sensitive to dithionite reduction within aLDs and thus provides a more accurate assessment of aLD surface accessibility and sample purity. By comparison, in the dithionite quenching assay, NBD-PC-labeled aLDs with the optimized protocol showed 81% quenching **(Fig. 5A)**.

**Figure 5.**
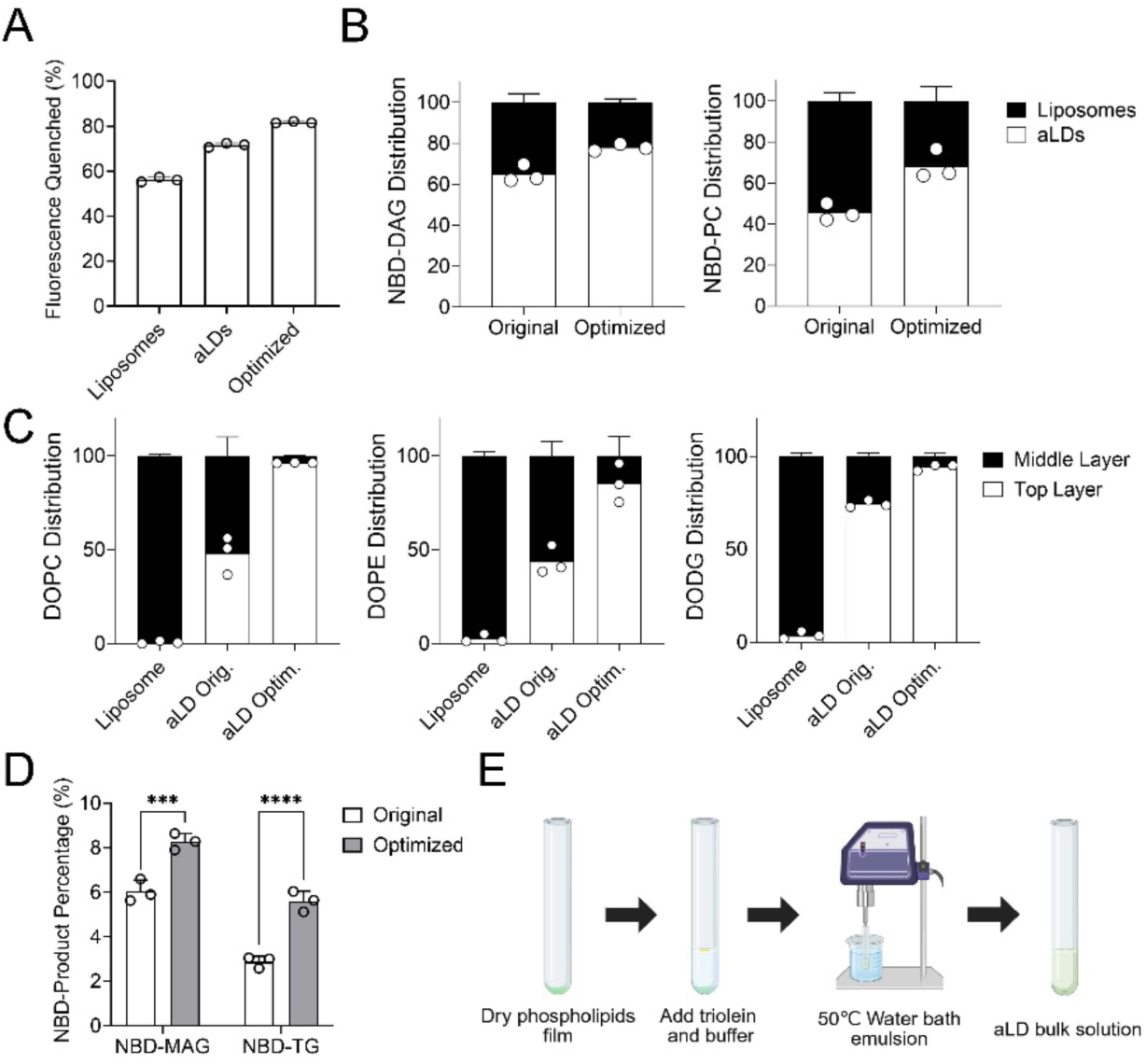
Enhanced aLD purity and lipid partitioning increases DDHD2 lipase and transacylase activities. **(A)** Comparison of the dithionite quenching percentage of NBD-PC labeled liposomes, aLDs generated with original protocol (original), and the optimized protocol (optimized). Mean ± SD is shown for each bar graph (n = 3). **(B)** Comparison of NBD-DAG or NBD-PC distribution in aLDs or liposomes after sucrose-gradient centrifugation. Mean ± SD is shown for each bar graph (n = 3). **(C)** Comparison of DOPC, DOPE, DODG (DAG) distribution in aLDs (top layer) or liposomes (middle layer) after sucrose-gradient centrifugation. Mean ± SD is shown for each bar graph (n = 3). **(D)** DDHD2 activity assay in original and optimized aLD system, quantified NBD-MAG and NBD-TAG products percentage. Mean ± SD is shown for each bar graph (n = 3). **(E)** Schematic of the optimized aLD preparation protocol. After drying phospholipids, triolein and buffer are added together and emulsified in a 50 °C water bath by microtip sonication.

### Quantitation of NBD and unlabeled lipid partitioning in aLDs versus liposomes

To further evaluate how liposome contamination affected the distribution of phospholipids and neutral lipids in the aLD system, we analyzed the top and middle fractions collected after sucrose gradient centrifugation by HPLC. aLDs were prepared with NBD-PC to label phospholipids or NBD-DAG to label neutral lipids. Using our original aLD preparation protocol, about 60% of the NBD-PC signal and roughly 40% of the NBD-DAG signal were found in the liposome fraction **(Fig. 5B)**. After applying the optimized preparation method, these proportions decreased to approximately 40% for NBD-PC and 20% for NBD-DAG **(Fig. 5B)**.

We also analyzed the distribution of unlabeled lipids by mass spectrometry (MS). As controls, we prepared a liposome-only sample, aLDs using the original protocol, and aLDs using the optimized protocol. After sucrose gradient centrifugation, the liposome-only sample showed 97% of DOPE, 99% of DOPC, and 96% of DO-DAG in the liposome fraction **(Fig. 5C)**. Using our original aLD protocol, we observed 56% DOPE, 52% DOPC, and 26% DO-DAG in the liposome fraction **(Fig. 5C)**. By comparison, the optimized aLD protocol showed 14% DOPC, 4% DOPE, and 6% DO-DAG in the liposome fraction **(Fig. 5C)**. This increased distribution of both NBD-lipids and unlabeled lipids within aLDs versus liposomes is consistent with increased aLD purity, thus providing a more suitable system for downstream in vitro protein assays.

### Optimized aLDs increase enzymatic activity in DDHD2

Next, we assessed the difference in lipase activity between the optimized and original aLD preparation protocols. DDHD2, recently described as a dual functional lipase that hydrolyzes diacylglycerol (DAG) to generate monoacylglycerol (MAG) and free fatty acids, and a transacylase that converts DAG to TAG, was chosen as an example. We prepared aLDs containing the same concentrations of phospholipids, NBD-DAG, and TAG using both the original and optimized methods, and incubated them with DDHD2. The optimized protocol gave rise to an approximately 2-fold increase in the generation of both NBD-MAG and NBD-TAG by DDHD2 compared with the original aLD protocol **(Fig. 5D, E)**. This indicates that the improved purity of the optimized aLDs increased the lipase and transacylase activities of DDHD2.

### Surface accessibility of neutral lipids within aLDs depends on hydrophobicity

Having optimized our aLD protocol, we next sought to assess the surface accessibility of different neutral lipids within aLDs using our biotin-streptavidin imaging system or dithionite quenching approach. Neutral lipids are stored within the oil core of lipid droplets, with TAG being the major component. Because neutral lipids differ in structure, acyl-chain number and acyl-chain length, their overall hydrophobicity can vary, which may influence their distribution within the droplet core. To test whether hydrophobicity affects the internal distribution of neutral lipids, we prepared aLDs containing different NBD-labeled neutral lipids, including NBD-DAG 10:0-10:0, NBD-DAG 18:0-20:4, NBD-cholesteryl ester, and NBD-TAG 18:0-18:0-18:0 **(Fig. 6A)**.

**Figure 6.**
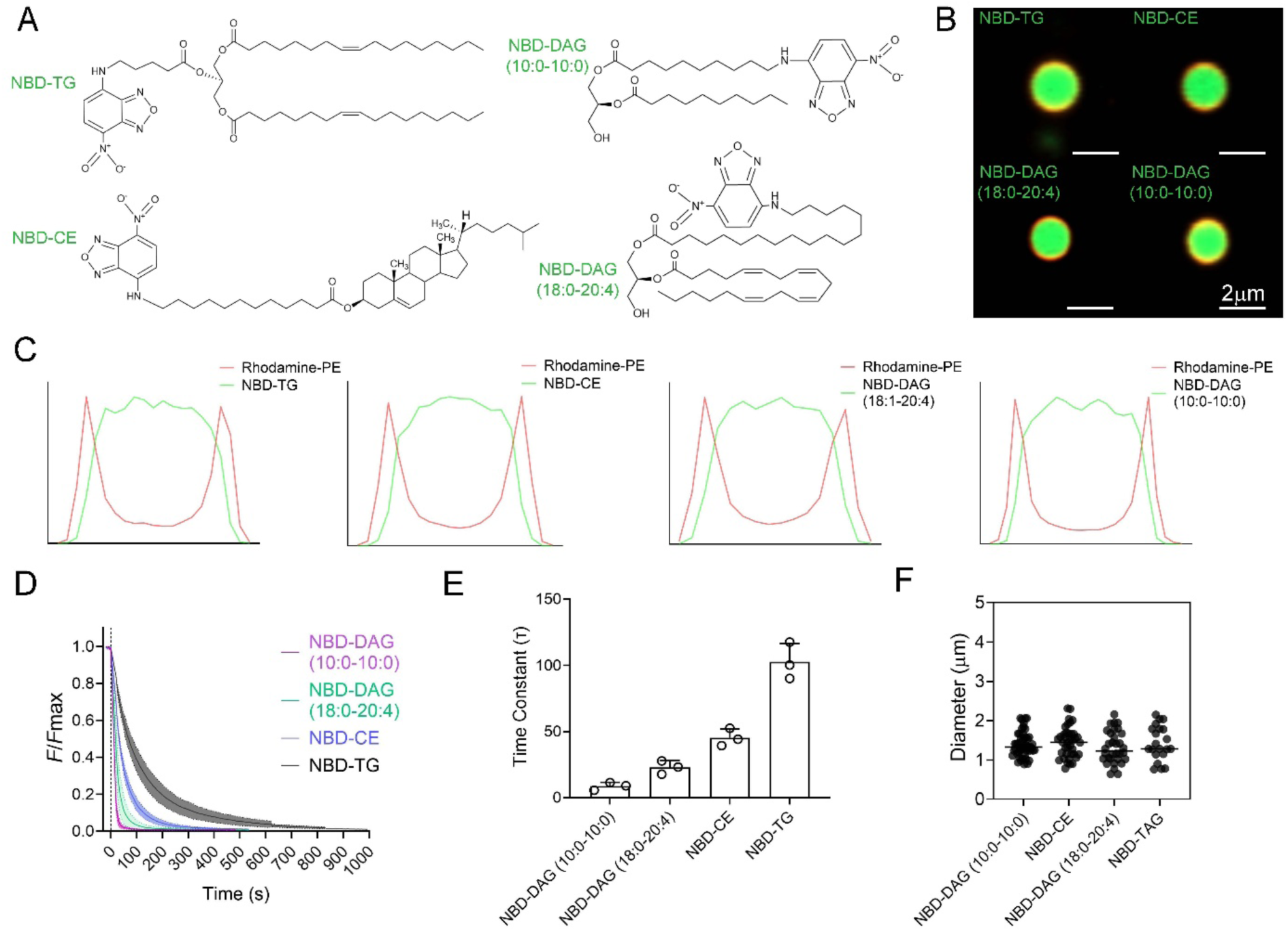
Hydrophobicity determines the surface accessibility of neutral lipids within aLDs. **(A)** Chemical structure for NBD-TG, NBD-DAG(10:0-10:0), NBD-CE, and NBD-DAG (18:0-20:4). **(B)** Fluorescence microscopy images of aLDs labeled with NBD-TG, NBD-CE, NBD-DAG (18:0-20:4), or NBD-DAG (10:0-10:0) (green) and rhodamine-PE (red). Scale bar = 2 μm. **(C)** Fluorescence intensity line functions with different NBD labeled neutral lipids. Quantified by Zen-Profile function. aLDs were centered by Z-stack imaging. **(D)** Normalized fluorescence intensity from dithionite quenching assay with aLDs containing either NBD-DAG (10:0-10:0), NBD-DAG (18:0-20:4), NBD-CE, or NBD-TG. (n=3) **(E)** Quantification of the time constant from dithionite quenching decay curve from (D). Mean ± SD is shown for each bar graph (n = 3). **(F)** Diameters of aLDs prepared with NBD-DAG (10:0/10:0), NBD-CE, NBD-DAG (18:0-20:4), or NBD-TAG were measured by confocal fluorescent microscopy. Each dot represents an individual measurement (n = 51, 39, 32, 21).

aLDs were immobilized using the biotin–streptavidin method and imaged with Airyscan confocal microscopy **(Fig. 6B)**. Z-stack images were collected for each droplet, and fluorescence distribution profiles were generated from slices across the droplet. The slice with the strongest rhodamine signal was used to define the droplet center **(Supplementary Fig. 2A)**. Mapping the fluorescence of the different NBD-labeled neutral lipids showed that all probes appeared similarly distributed within the droplet core, and no significant differences were detected visually across lipids with different hydrophobicities **(Fig. 6C)**.

Since several variables, including resolution and FRET between NBD and rhodamine fluorophores, might limit the use of microscopy to detect differences in solvent exposure when most of an NBD-labeled lipid is within the core of the aLD, we next measured the accessibility of these NBD-labeled neutral lipids using dithionite quenching. Dithionite can reduce NBD-neutral lipid molecules that are surface exposed near the phospholipid monolayer. This would allow quenching kinetics to report on how readily each probe interacts with the droplet surface.

The quenching results clearly showed a dependence on hydrophobicity **(Fig. 6D)**. NBD-DAG (10:0-10:0) was quenched rapidly with a time constant of ∼10 s, whereas NBD-TAG showed much slower quenching with a time constant of ∼109 s. NBD-DAG with longer acyl chains (18:0-20:4) and NBD-CE had intermediate time constants of 20 s and 45 s, respectively **(Fig. 6E)**. While we could not control aLD size, which could affect the observed quenching rates, we did not observe a dependence of aLD diameter on the NBD neutral lipid species (**Supplementary Fig. 2B)**, with all aLDs having similar size distributions, ranging mainly between 1–2 µm (**Fig. 6F**). Taken together, these data indicate that the accessibility of neutral lipids to the droplet surface decreases with increasing hydrophobicity, which we were unable to detect using a canonical microscopy approach.

## DISCUSSION

In this study, we demonstrate dithionite quenching of NBD-labeled lipids as a sensitive and quantitative approach for assessing aLD purity and neutral lipid accessibility. While fluorescence microscopy and electron microscopy are effective for confirming overall droplet morphology, these methods do not readily detect contaminating membrane structures, such as liposomes, or provide information about probe accessibility at the droplet surface. By exploiting the selective reduction of NBD fluorophores exposed to the aqueous phase, dithionite quenching enabled us to distinguish monolayer-coated lipid droplets from bilayer membrane structures and to quantify the extent of liposome contamination in bulk aLD preparations. Importantly, this assay can be applied rapidly and without extensive sample manipulation, making it well suited for routine quality control of aLD preparations.

Beyond their utility for purity assessment, the optimized bulk aLD preparations provide an ideal platform for biochemical assays of enzymes that function at lipid droplets. The enhanced DDHD2 activity we observed with our optimized aLD protocol suggests that our initial characterization of DDHD2 (46) may have underestimated its activity on droplets, given a substantial fraction of the fluorescent substrate partitions into contaminating liposomes. Optimization of aLD purity is thus recommended for any biochemical experiments that rely on bulk solution experiments.

Dithionite quenching also revealed insights into lipid organization within the droplet monolayer. Although the NBD group is somewhat polar, and our kinetic experiments could not be carried out with unlabeled lipids, the differences in quenching kinetics demonstrated that neutral lipid accessibility within artificial lipid droplets depends on lipid hydrophobicity. Less hydrophobic lipids exhibited faster quenching kinetics, which suggests that more hydrophobic lipids have reduced exposure to the phospholipid monolayer interface. This differential accessibility may have implications for lipases that access substrates at the monolayer interface. For example, during lipolysis, conversion of TAG to DAG and then MAG, generates progressively less hydrophobic intermediates that may have greater access to the droplet surface. This could facilitate continued substrate processing by lipases. In future applications, this approach could be extended to evaluate how lipid composition or protein binding alters the properties of neutral lipids in aLDs.

A current limitation of the aLD preparation method is the inability to control aLD size. In principle, the droplet diameter can be estimated from the available phospholipid surface area relative to the neutral lipid core volume at different phospholipid-to-triacylglycerol (PL: TAG) ratios. Adjusting the PL: TAG ratio should therefore enable predictable modulation of droplet size. However, in our experiments, the measured droplet diameters did not consistently match the calculated values. This discrepancy likely reflects the incomplete incorporation of phospholipids into the droplet monolayer and the incorporation of these phospholipids into liposomes. As a result, the effective phospholipid coverage of the aLD monolayer may differ from theoretical assumptions. While the current preparation yields reproducible average droplet populations of ∼1μm that are suitable for comparative assays, future refinement is needed to accurately predict and control the size of aLDs for bulk biochemical studies.

## METHODS

### Lipids and reagents

Dioleoyl-phosphatidylcholine (DOPC, Cat. #850375), Dioleoyl-phosphatidylethanolamine (DOPE, Cat. #850725), 18:1-12:0 NBD PC (NBD-PC, Cat. #810133),18:1 NBD PE (NBD-PE, Cat. #810145), 18:1 Biotinyl Cap PE (Biotin-PE, Cat. #870273), and 18:1 Liss Rhod PE (Rh-PE, Cat. #810150) were purchased from Avanti Polar Lipids. 1-NBD-decanoyl-2-decanoyl-sn-Glycerol (NBD-DAG, Cat. #9000341), 1-NBD-Stearoyl-2-Arachidonoyl-sn-glycerol (NBD-SAG, Cat #10011300), and Sodium hydrosulfite (Sodium Dithionite, Cat. #71699) were purchased from Sigma Aldrich. 3-dodecanoyl-NBD Cholesterol (NBD-CE, Cat. #13220), 1,2,3-Trioleoyl Glycerol (TG, Cat. #26871) were purchased from Cayman Chemical. 1,3-diolein, 2-NBD-X ester (NBD-TG, Cat. #6285) was purchased from Setareh Biotech.

### Initial Protocol to Prepare ALDs

In the prepared aLDs, phospholipids (PLs) and triglycerides (TAG) accounted for 5.2% and 94.8% of the total lipids, respectively, with a PL:TG ratio of 1:18. In the initial protocol to prepare aLDs, 15.67 μL of DOPC (stock conc. 31.8 mM) and 3.71 μL of DOPE (stock conc. 33.6 mM) in chloroform were mixed in a glass tube and dried under a stream of nitrogen gas. After, 10 μL of TG (stock conc. 1124 mM) was added. The PL:TG mixture was then sonicated in a water bath sonicator (BRANSON Ultrasonic Cleaner) at RT for 20 min to solubilize the PLs in the TG oil. After premixing, 490 μL of 50 mM HEPES, 50 mM NaCl, pH 8 aqueous buffer (HN Buffer) was added, gently vortexed for 30 s at a minimal setting, and underwent a second round of water bath sonication at RT for 20 min. At this point, the solution was slightly turbid, and ultrasonic emulsification was performed using tip sonication with a Fisherbrand™ Model 120 Sonic Dismembrator (catalog# FB120110) with a 1/8 in. (3.2mm) Titanium Probe (13.8cm x 1.3cm, Titanium Alloy, catalog# FB4422) with the glass tube submerged in a RT water bath. Initial tip sonication settings were: 35% amplitude, with 2 s on/ off cycles for 5 min. A schematic of this initial protocol is shown in Supplementary Fig. 1. The micro-tip used in this study showed moderate pitting consistent with normal cavitation wear, with no cracking or chipping of the tip edge or shaft **(Supplementary Fig. 3)**.

### Optimized Protocol to Prepare ALDs

The composition of the phospholipid components was the same as in the original protocol. After drying under a stream of nitrogen, 10 μL of TG (18:1/18:1/18:1) (1124 mM) was added, and 490 μL of buffer (50 mM HEPES, 50 mM NaCl, pH 8) was added. The mixture was gently vortexed for 30 s. After premixing, the mixture was ultrasonically emulsified using a Fisherbrand™ Model 120 Sonic Dismembrator at 40% amplification, 2 s on / 2 s off cycles, in a 50°C water bath for 5 min.

### Liposome Preparation

The composition of the phospholipid components was the same as in the aLD protocol. After drying under a stream of nitrogen, 500 μL of buffer (50 mM HEPES, 50 mM NaCl, pH 8) was added. The mixture was gently vortexed for 30 s. After premixing, the mixture was ultrasonically emulsified using a Fisherbrand™ Model 120 Sonic Dismembrator at 40% amplification, 2 s on / 2 s off cycles for 1 min, sonicating two cycles.

### Dithionite Quenching Assay

174 mg (MW, 174.1 g/mol) of sodium dithionite powder was freshly dissolved in 1 mL of Tris-HCl buffer (1M Tris, pH 10.0) to produce a 1 M solution. Liposomes or aLDs, which were freshly prepared within 2 h, were diluted 100-fold in HN buffer and 100 μL were added to each well of a Costar® 96-Well Black Polystyrene Plate. Fluorescence intensity of the samples was measured every 5 s for 50 s using a Molecular Devices SpectraMax M2 Multilabel Microplate Reader in SoftMax Pro 7.1 in FL read mode and kinetic read type, using excitation = 479 nm, emission = 530 nm. Subsequently, addition of 2 μL of freshly prepared 1M sodium dithionite solution was added to each sample, and mixed quickly. The real-time fluorescence intensity was measured using the same protocol until the fluorescence intensity stabilized, which typically occurred after 5 min. Finally, 10 μL of a 1% w/v Triton-X 100 solution was added and fluorescence readings were measured for 1 min until the fluorescence intensity stabilized.

### Biotin-Streptavidin Anchoring

Glass slides (Cat #632010, Carolina) were coated with 50 μL of a 5% BSA solution, spread evenly, and air-dried at RT for 30 min. Coverslips (Cat #48366-045, VWR) were coated with 20 μL of 1% biotin-BSA (Cat #A8549, Sigma Aldrich), in the same buffer as aLDs, were spread evenly for 30 min to form a thin film. Coverslips were rinsed once with HN buffer and then incubated with 30 μL of a 1% streptavidin solution (Cat #434301,Thermo Fisher) for 10 min at RT, and later rinsed 3 times with 200 μL aLD buffer. Diluted 10-fold aLD suspensions containing 1 mol% biotin-PE were applied to the BSA-precoated glass slides (15 μL per slide), and coverslips were mounted carefully to avoid compression of the droplets. Slides were sealed with nail polish.

### Sucrose Gradient Separation

aLDs were prepared in a 30% w/v sucrose solution in HN buffer (30% sucrose w/v, 50 mM HEPES, 50 mM NaCl, pH 8). Prior to centrifugation, the following solutions were added sequentially to an 11×34 mm Polycarbonate Centrifuge Tube (Beckman Coulter, Cat #343778): 1) 400 μL of HN buffer, 2) 200 μL of a two times dilution of the bulk aLD solution, and 3) 200 μL of a 30% w/v sucrose solution in HN buffer (30% w/v). Centrifugation was performed using an Optima MAX-XP Ultracentrifuge TLS-55 Swinging-Bucket Rotor at 35,000 xg at 4°C for 30 min. Following centrifugation, fractions were collected from the bottom of the centrifuge tube using a blunt-tip syringe. The bottom 150 μL fraction was first withdrawn and discarded. Using a new syringe, the next 350 μL fraction was collected and transferred to a clean centrifuge tube. The syringe was then replaced again, and the remaining 300 μL fraction was collected into a separate centrifuge tube. The middle fraction contained the interface between the 0% and 30% sucrose layers, where contaminating liposomes were enriched after centrifugation. The final top fraction consisted of the 0% sucrose layer and the floating lipid droplet layer, representing the purified aLD fraction.

### NBD Labeled Lipid Distribution

aLDs were separately prepared with NBD-DAG and NBD-PC in a 30% w/v sucrose solution in HN buffer (30% sucrose w/v, 50 mM HEPES, 50 mM NaCl, pH 8). After sucrose gradient separation, the middle (liposome) and top (aLD) fractions were collected as described in the “Sucrose Gradient Separation” section above, and the lipids in each fraction were extracted by addition of a methanol:chloroform (1:1, v:v) solution and vortexed vigorously to mix. The mixture was centrifuged at 3,000×*g* for 5 min, the lower organic phase was extracted using a glass Hamilton syringe and transferred to a new glass tube. After the organic phase was dried under nitrogen gas for 30 min and resuspended in HPLC Buffer B containing 100% methanol, 1 mM ammonium formate and 0.2% formic acid (v/v). The solution was centrifuged at 3,000×*g* for 5 min and the supernatant was transferred to an HPLC vial with inserts.

### Unlabeled Lipid Distribution

aLDs were separately prepared with DOPC, DOPE, and DO-DAG in a 30% w/v sucrose solution in HN buffer (30% sucrose w/v, 50 mM HEPES, 50 mM NaCl, pH 8). Liposome controls were prepared using the same concentration of DOPC, DOPE, and DAG as in the aLDs but without triolein. After sucrose gradient separation, the middle (liposome) and top (aLD) fractions were collected as described above, and the lipids in each fraction were extracted by addition of a methanol:chloroform (1:1, v:v) solution and vortexed vigorously to mix. The mixture was centrifuged at 3,000×*g* for 5 min, the lower organic phase was extracted using a glass Hamilton syringe and transferred to a new glass tube. The organic phase was dried under nitrogen gas for 30 min and resuspended in 100% methanol. The solution was centrifuged at 3,000×g for 5 min, the supernatant was transferred to a fresh tube, and 50 μL of PC 13:0/13:0 internal standard. Samples were dried under nitrogen gas and dissolved in 300 μL mobile solvent B (1% H2O/9% acetonitrile/90% isopropanol (v/v/v), 10 mM ammonium formate). Samples were diluted 1:10 and injected in an Agilent 6490 coupled to a 1290 series HPLC system with ZORBAX RRHD Eclipse Plus C18 1.8 μm column (Agilent catalog # 959758-902). Parameters were set as peak width 0.02 min, fragmentor 250 V, positive polarity, drying gas 10 L/min, nebulizer 20 psi, drying gas temperature 350 °C. Flow: 0.4 mL/min; Mobile solvent A: 50% H2O/30% acetonitrile/20% isopropanol (v/v/v),10 mM ammonium formate; Start with 90% solvent A, 10% solvent B; Gradient: 0 to 2.7 min 10 to 45% solvent B, 2.7 to 2.8 min 45 to 53% solvent B, 2.8 to 9 min 53 to 65% solvent B, 9 to 9.1 min 65 to 89% solvent B, 9.1 to 11 min 89 to 92% solvent B, 11 to 13.9 min 92 to 100% solvent B, 13.9 to 14.9 min 100 to 10% solvent B. Column was equilibrated for 3 min before the next injection. The MRM table for the detected species was: PC(18:1/18:1) 786.6/184.1, PE(18:1/18:1) 744.6/603.5, DG(18:1/18:1) 638.5/339.3 and PC(13:0/13/0) 650.5/184.1. The area under the curve was calculated using MassHunter Software. All compounds were normalized using the internal standard.

### In Vitro DDHD2 Activity Assay by HPLC

Full-length recombinant human DDHD2 was expressed and purified as previously described (36). aLDs were generated separately using the original and optimized protocols, with calculated amounts of DOPC, DOPE, and NBD-DAG (10:0-10:0) in an 80:20:2 molar ratio, yielding final concentrations of 1 mM DOPC, 0.25 mM DOPE, and 25 μM NBD-DAG (10:0-10:0) substrate. The prepared substrate containing aLDs and protein were mixed at a 1:1 volume ratio and incubated for 1 h at 37 °C. The reaction was terminated by addition of a methanol:chloroform (1:1, v:v) solution and vortexed vigorously to mix. The mixture was centrifuged at 3,000×*g* for 5 min, the lower organic phase was extracted using a glass Hamilton syringe and transferred to a new glass tube. After, the organic phase was dried under nitrogen gas for 30 min and resuspended in HPLC Buffer B containing 100% methanol, 1 mM ammonium formate and 0.2% formic acid (v/v). The solution was centrifuged at 3,000×*g* for 5 min and the supernatant was transferred to an HPLC vial with inserts. The above samples were analyzed in an Agilent HPLC system with the Spectra 3 μm C8SR column (Catalog # S-3C8SR-FJ) under the following conditions:Injection volume: 10 μL; Flow: 0.5 mL/min; Mobile phase A: Fisher Water Optima LC/MS, 1 mM ammonium formate, 0.2% formic acid (v/v); Mobile phase B: Fisher Methanol Optima, 1 mM ammonium formate, 0.2% formic acid (v/v); Gradient: 0 to 1 min 50 to 80% Buffer B, 1 to 8 min 80 to 98% Buffer B, 8 to 11 min 98% Buffer B, then 11 to 16 min 80% Buffer B. The fluorescence detector was set to scan excitation wavelength 470 nm and emission wavelength 530 nm according to the properties of the NBD fluorophore.

### Fluorescence Microscopy of ALDs

Biotin-Streptavidin-immobilized aLD slides were imaged inverted using a Zeiss LSM 980 Airyscan 2 NLO Two-Photon Confocal Microscope, using a 100x oil objective and ZEN software for high-resolution image capture and image analysis. Lipid droplets labeled with NBD-neutral lipids (0.1 mol% total lipids) and Rhodamine-PE (1 mol% PLs) were imaged using the Airyscan TaRFP (excitation/emission/effective NA 558/583/1.4) and Airyscan EGFP (excitation/emission/effective NA 488/509/1.4). For size quantification, z-stacks were acquired at 0.17 µm intervals. Using the profile analysis tool in the ZEN software, the cross-section with the maximum spacing and intensity of the Rhodamine-PE fluorescence peaks was selected as the equatorial plane of each droplet. The peak-to-peak distance of the Rhodamine-PE intensity profile at this plane was recorded as the droplet diameter and used to compare aLD size across preparations containing different NBD-labeled neutral lipids.

### Statistical Analysis

Statistical analysis was performed using one-way ANOVA followed by Tukey’s post hoc test for multiple comparisons. Data are presented as mean ± standard deviation (SD). A *p*-value of less than 0.05 was considered statistically significant.

## ACKNOWLEDGMENTS

We wish to acknowledge the Stony Brook Cancer Center Biological Mass Spectrometry Shared Resource for expert assistance with lipidomics analysis. We thank Dr. Aaron Neiman for sharing access to his fluorescence microscope, and Kai Zhang and Dr. Jae Sook for their guidance in its use. We also thank Dr. Shinako Kakuda for her guidance on the sodium dithionite quenching assay. This work was supported by the NIH Grants GM128666 (M.V.A.) and GM122493 (E.L.), a Sloan Research Fellowship (M.V.A.), startup funds from the Stony Brook Cancer Center (D.C.), and the Agence Nationale de la Recherche, ANR-21-CE13-0014-LIPDROPER (A.R.T.).

## REFERENCES

1. T. Fujimoto, R. G. Parton, Not just fat: the structure and function of the lipid droplet. Cold Spring Harb Perspect Biol 3 (2011).

2. T. C. Walther, R. V. Farese, Jr., Lipid droplets and cellular lipid metabolism. Annu Rev Biochem 81, 687–714 (2012).

3. K. Tauchi-Sato, S. Ozeki, T. Houjou, R. Taguchi, T. Fujimoto, The surface of lipid droplets is a phospholipid monolayer with a unique Fatty Acid composition. J Biol Chem 277, 44507–44512 (2002).

4. C. Francois-Martin, F. Pincet, Actual fusion efficiency in the lipid mixing assay - Comparison between nanodiscs and liposomes. Sci Rep 7, 43860 (2017).

5. Q. Lin, E. London, Preparation of artificial plasma membrane mimicking vesicles with lipid asymmetry. PLoS One 9, e87903 (2014).

6. A. R. Thiam, E. Ikonen, Lipid Droplet Nucleation. Trends Cell Biol 31, 108–118 (2021).

7. T. P. Eva Jarc, Droplets and the Management of Cellular Stress. The Yale journal of biology and medicine 92, 435–452 (2019).

8. M. F. Renne, H. Hariri, Lipid Droplet-Organelle Contact Sites as Hubs for Fatty Acid Metabolism, Trafficking, and Metabolic Channeling. Frontiers in Cell and Developmental Biology 9 (2021).

9. M. Doktorova et al., Preparation of asymmetric phospholipid vesicles for use as cell membrane models. Nat Protoc 13, 2086–2101 (2018).

10. A. R. Thiam, R. V. Farese, Jr., T. C. Walther, The biophysics and cell biology of lipid droplets. Nat Rev Mol Cell Biol 14, 775–786 (2013).

11. A. Chorlay, A. R. Thiam, An Asymmetry in Monolayer Tension Regulates Lipid Droplet Budding Direction. Biophys J 114, 631–640 (2018).

12. A. Chorlay, L. Foret, A. R. Thiam, Origin of gradients in lipid density and surface tension between connected lipid droplet and bilayer. Biophys J 120, 5491–5503 (2021).

13. A. R. Althaher, An Overview of Hormone-Sensitive Lipase (HSL). ScientificWorldJournal 2022, 1964684 (2022).

14. F. Deslandes, A. R. Thiam, L. Foret, Lipid Droplets Can Spontaneously Bud Off from a Symmetric Bilayer. Biophys J 113, 15–18 (2017).

15. A. Santinho, A. Chorlay, L. Foret, A. R. Thiam, Fat inclusions strongly alter membrane mechanics. Biophys J 120, 607–617 (2021).

16. K. Ben M’barek et al., ER Membrane Phospholipids and Surface Tension Control Cellular Lipid Droplet Formation. Dev Cell 41, 591–604 e597 (2017).

17. P. Zhao et al., Artificial Lipid Droplets: Novel Effective Biomaterials to Protect Cells against Oxidative Stress and Lipotoxicity. Nanomaterials (Basel) 12 (2022).

18. A. Chorlay, A. Santinho, A. R. Thiam, Making Droplet-Embedded Vesicles to Model Cellular Lipid Droplets. STAR Protoc 1, 100116 (2020).

19. Z. Telikani et al., The phospholipid composition of artificial lipid droplets enhances their deliverability and facilitates a broad Biodistribution in vivo and in vitro. Acta Biomater 200, 520–535 (2025).

20. D. L. Brasaemle, V. Subramanian, A. Garcia, A. Marcinkiewicz, A. Rothenberg, Perilipin A and the control of triacylglycerol metabolism. Mol Cell Biochem 326, 15–21 (2009).

21. A. Lass, R. Zimmermann, M. Oberer, R. Zechner, Lipolysis - a highly regulated multi-enzyme complex mediates the catabolism of cellular fat stores. Prog Lipid Res 50, 14–27 (2011).

22. T. C. Walther, R. V. Farese, Jr., The life of lipid droplets. Biochim Biophys Acta 1791, 459–466 (2009).

23. A. R. Thiam et al., COPI buds 60-nm lipid droplets from reconstituted water-phospholipid-triacylglyceride interfaces, suggesting a tension clamp function. Proc Natl Acad Sci U S A 110, 13244–13249 (2013).

24. S. A. Gandhi et al., Methods for making and observing model lipid droplets. Cell Rep Methods 4, 100774 (2024).

25. A. Chorlay, A. R. Thiam, Neutral lipids regulate amphipathic helix affinity for model lipid droplets. J Cell Biol 219 (2020).

26. A. Chattopadhyay, E. London, Parallax method for direct measurement of membrane penetration depth utilizing fluorescence quenching by spin-labeled phospholipids. Biochemistry 26, 39–45 (1987).

27. T. C. Walther, J. Chung, R. V. Farese, Jr., Lipid Droplet Biogenesis. Annu Rev Cell Dev Biol 33, 491–510 (2017).

28. Y. Wang et al., Construction of Nanodroplet/Adiposome and Artificial Lipid Droplets. ACS Nano 10, 3312–3322 (2016).

29. X. Ma et al., Validating an artificial organelle: Studies of lipid droplet-specific proteins on adiposome platform. iScience 24, 102834 (2021).

30. Z. Zhi et al., Protocol for using artificial lipid droplets to study the binding affinity of lipid droplet-associated proteins. STAR Protoc 3, 101214 (2022).

31. S. F. Shimobayashi, Y. Ohsaki, Universal phase behaviors of intracellular lipid droplets. Proc Natl Acad Sci U S A 116, 25440–25445 (2019).

32. R. Dhiman, S. Caesar, A. R. Thiam, B. Schrul, Mechanisms of protein targeting to lipid droplets: A unified cell biological and biophysical perspective. Semin Cell Dev Biol 108, 4–13 (2020).

33. F. S. Abrams, E. London, Extension of the parallax analysis of membrane penetration depth to the polar region of model membranes: Use of fluorescence quenching by a spin-label attached to the phospholipid polar headgroup. Biochemistry 32, 10826–10831 (1993).

34. C. A. a. J. W. Nichols, Dithionite Quenching Rate Measurement of the Inside-Outside Membrane Bilayer Distribution of 7-Nitrobenz-2-oxa-1,3-diazol-4-yl-Labeled Phospholipids. Biochemistry 37 (1998).

35. K. Bersuker et al., A Proximity Labeling Strategy Provides Insights into the Composition and Dynamics of Lipid Droplet Proteomes. Dev Cell 44, 97–112 e117 (2018).

36. L. Wu et al., DDHD2 possesses both lipase and transacylase capacities that remodel triglyceride acyl chains. Proc Natl Acad Sci U S A 122, e2500527122 (2025).

37. J. L. Korfhage et al., ATG2A-mediated bridge-like lipid transport regulates lipid droplet accumulation. bioRxiv 10.1101/2023.08.14.553257 (2023).

38. C. Angeletti, J. W. Nichols, Dithionite Quenching Rate Measurement of the Inside−Outside Membrane Bilayer Distribution of 7-Nitrobenz-2-oxa-1,3-diazol-4-yl-Labeled Phospholipids. Biochemistry 37, 15114–15119 (1998).

39. P. T. O’Neil, A Limitation of Using Dithionite Quenching to Determine the Topology of Membrane-inserted Proteins. J Membr Biol 255, 123–127 (2022).

40. M. J. Moreno, L. M. Estronca, W. L. Vaz, Translocation of phospholipids and dithionite permeability in liquid-ordered and liquid-disordered membranes. Biophys J 91, 873–881 (2006).

41. R. Song, Y. Lin, Z. Li, Ultrasonic-assisted preparation of eucalyptus oil nanoemulsion: Process optimization, in vitro digestive stability, and anti-Escherichia coli activity. Ultrason Sonochem 82, 105904 (2022).

42. S. Manley, M. R. Horton, S. Lecszynski, A. P. Gast, Sorting of streptavidin protein coats on phase-separating model membranes. Biophys J 95, 2301–2307 (2008).

43. D. H. Murray, L. K. Tamm, V. Kiessling, Supported double membranes. J Struct Biol 168, 183–189 (2009).

44. S. J. Schmidt A, Bayerl T, Sackmann E, Knoll W, Streptavidin binding to biotinylated lipid layers on solid supports. A neutron reflection and surface plasmon optical study. Biophysical journal 63, 1385–1392 (1992).

45. C. Joo, T. Ha, Preparing sample chambers for single-molecule FRET. Cold Spring Harb Protoc 2012, 1104–1108 (2012).

46. L. Wu et al., DDHD2 possesses both lipase and transacylase capacities that remodel triglyceride acyl chains. bioRxiv 10.1101/2025.09.22.677815 (2025).

