## Supplemental Information for "Dithionite quenching of NBD-labeled lipids reveals artificial lipid droplet purity and neutral lipid surface accessibility"

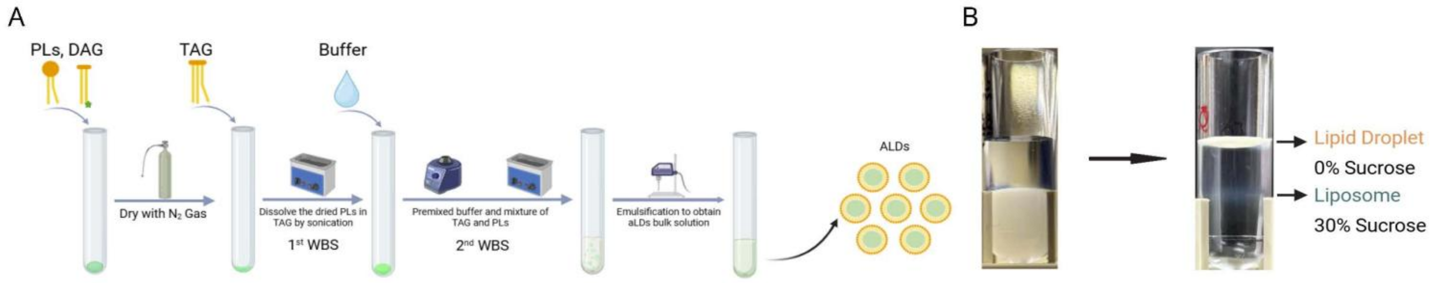


**Supplementary Figure 1. Artificial lipid droplet preparation and sucrose centrifugation.**

**(A)** Schematic of the original artificial lipid droplet preparation protocol. Phospholipids and fluorescently labeled neutral lipids were dried under nitrogen in a glass test tube to form a thin lipid film. Triolein was added and mixed by water-bath sonication (1$^{st}$ WBS). HN buffer was then introduced, followed by brief vortexing and a second water-bath sonication (2$^{nd}$ WBS) step. The mixture was finally emulsified by microtip sonication to generate the bulk artificial lipid droplet solution. **(B)** Sucrose centrifugation leads to a purified aLD layer (top), and liposome layer (middle).

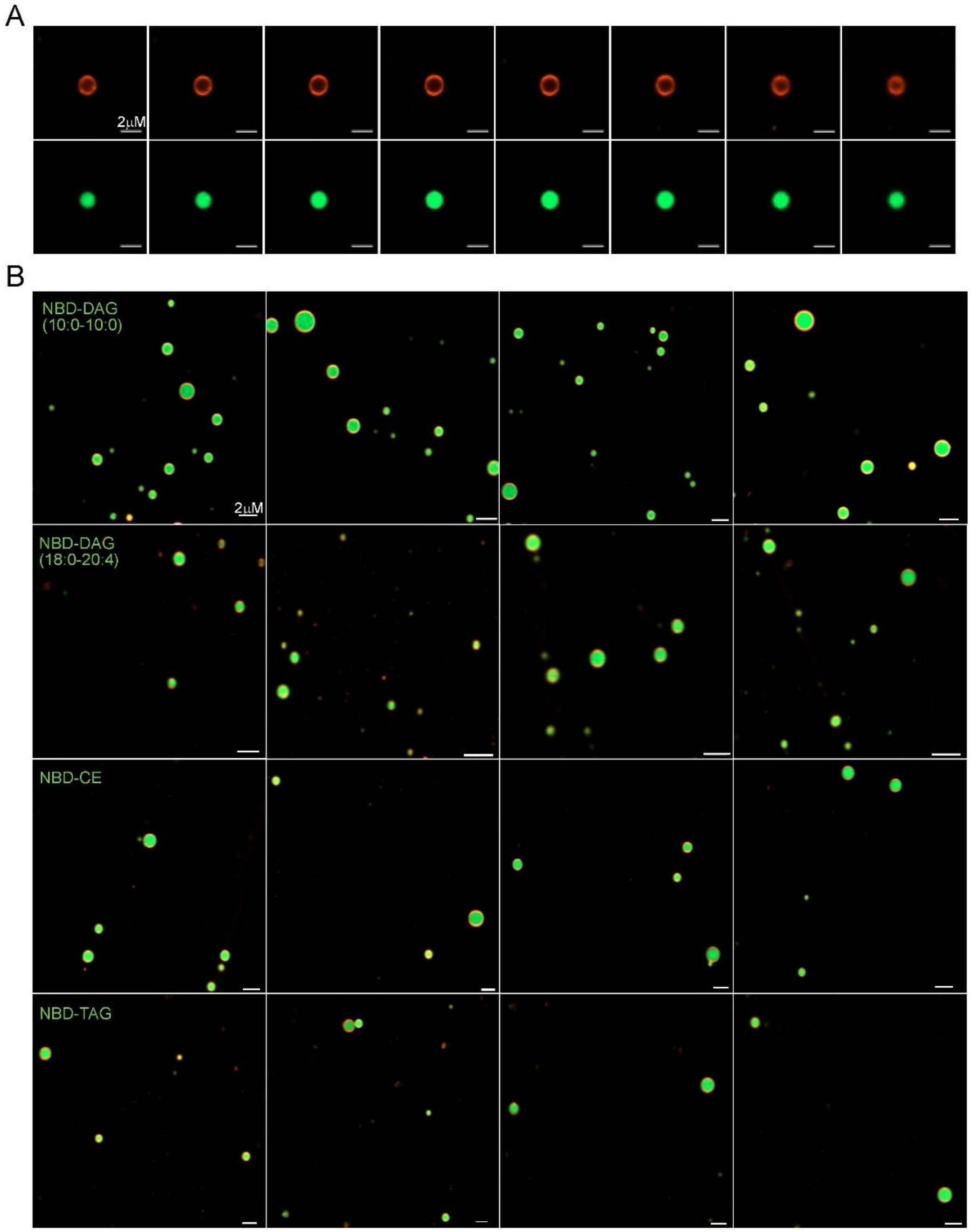


**Supplementary Figure 2. Z-stack image for a single aLD.**

**(A)** Z-stack confocal fluorescence imaging of the representative aLD shown in Fig. 6B. The droplet contains NBD-DAG (10:0–10:0) as the fluorescent neutral lipid probe. Rhodamine-PE, shown in red, marks the phospholipid monolayer, while NBD-DAG, shown in green, reports the distribution of the labeled neutral lipid. Optical sections were acquired across the z-axis of the droplet at 0.17 μm intervals between adjacent planes. Scale bar = 2 μm. **(B)** Confocal fluorescence imaging for artificial lipid droplets labeled with NBD-DAG (10:0-10:0), NBD-DAG (18:0-20:4), NBD-CE, or NBD-TAG (green) and rhodamine-PE (red). Scale bar = 2 μm.

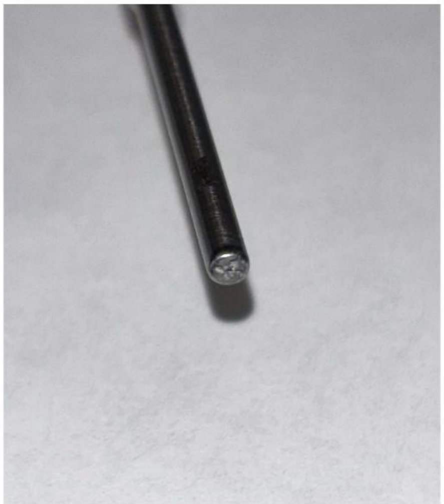

**Supplementary Figure 3. Sonicator Micro-tip.**

Picture of the titanium probe and micro-tip used in this study: a Fisher scientific 1/8 in. (3.2mm) Probe (13.8cm x 1.3cm) for processing 0.5-15 mL volumes (catalog# FB4422).
